# Inducible nitric oxide synthase (iNOS) regulates skin eschar lesions, bacterial persistence, and inflammatory resolution in mouse models of scrub typhus

**DOI:** 10.64898/2026.04.15.718641

**Authors:** Yixuan Zhou, Lihai Gao, Ryan H. Cho, Judy Ly, Hui Wang, Hema P. Narra, Kun-Hsien Tsai, Lynn Soong, Yuejin Liang

## Abstract

*Orientia tsutsugamushi* (*Ot*) is an obligately intracellular bacterium that causes scrub typhus, a potentially severe infectious disease characterized by systemic inflammation and multiorgan dysfunction. We recently reported a protective role for IFN-γ signaling in host defense against *Ot* infection; however, the underlying mechanisms remain obscure. Inducible nitric oxide synthase (iNOS, encoded by *Nos2*) is a key antimicrobial effector induced downstream of IFN-γ signaling. Here, we used transgenic mouse models to further investigate the biological functions of iNOS. We first revealed the requirement of iNOS for the restriction of *Ot* growth in cultured bone marrow-derived macrophages. Using an intradermal mouse model, we found that while tissues of *Nos2*^−/−^ and wild-type mice exhibited comparable bacterial burdens during acute infection phases, *Nos2*^−/−^ mice developed eschar-like lesions similar to those observed in *Ifngr1*^−/−^ mice, indicating a critical role for the IFN-γ/iNOS axis in regulating skin pathology in scrub typhus. Notably, *Nos2*^−/−^ mice displayed impaired bacterial clearance during the recovery phase (day 42), with persistent bacterial burdens in multiple organs accompanied by sustained immune activation and elevated inflammatory responses. Histopathological and biochemical analyses further revealed increased tissue damage and dysregulated physiological homeostasis in *Nos2*^−/−^ mice during recovery. Mechanistically, iNOS deficiency resulted in heightened myeloid cell activation and prolonged expression of proinflammatory mediators, suggesting a dual contribution of iNOS in both antimicrobial defense and inflammation resolution. Collectively, these findings provide new insight into IFN-γ-mediated defense mechanisms and imply the distinct roles of iNOS during different stages of scrub typhus.

**Author summary:** Scrub typhus is a potentially severe infectious disease caused by the bacterium *Orientia tsutsugamushi* (*Ot*), which is transmitted to humans through the bite of infected mites. Despite its global impact and expanding geographic distribution, the immune mechanisms that protect against this infection remain incompletely understood. In this study, we examined the role of inducible nitric oxide synthase (iNOS), an immune effector molecule that helps the host control infection. Using mouse models, we found that iNOS plays dual and stage-specific roles during *Ot* infection. Mice lacking iNOS developed dysregulated immune homeostasis during acute infection and exhibited skin lesions resembling the eschars observed in some patients with scrub typhus. In addition, these mice showed delayed bacterial clearance, prolonged inflammation, and increased tissue damage during the recovery phase. Our findings indicate that iNOS contributes not only to host antimicrobial defense but also to the control of excessive inflammation following infection. These results provide new insight into host defense mechanisms in scrub typhus and may help inform future therapeutic or preventive strategies.

## Introduction

*Orientia tsutsugamushi* (*Ot*) is an obligately intracellular bacterium and the etiological agent of scrub typhus, an acute febrile illness that can progress to severe systemic inflammation and multiorgan dysfunction^1–3^. Scrub typhus was historically thought to be confined to the so-called “tsutsugamushi triangle” in the Asia-Pacific region; however, new cases have now been reported well beyond this area, including the Middle East and South America^4^. It is estimated that there are more than one million new cases annually^5^. A meta-analysis reported a pooled seroprevalence of 10.73% among healthy individuals and 22.58% among febrile patients in endemic regions^6^. More recently, molecular detection of *Ot* species has been documented in free-living *Eutrombicula* chiggers in North Carolina, USA^7^. Consistently, in this region, antibodies reactive to *Ot* have also been detected in individuals presenting with skin eschars^8^, the most pathognomonic clinical sign of scrub typhus, suggesting that scrub typhus may be underrecognized in the United States. The clinical presentation typically involves abrupt onset of fever with chills, along with nonspecific symptoms including headache, myalgia, sweating, and vomiting^9^. Although scrub typhus is treatable with antibiotics such as doxycycline and azithromycin, diagnostic delay and antibiotic therapy failure can result in severe disease outcomes often associated with increased morbidity and mortality^10–12^. Currently, no licensed vaccine is available, largely due to the extensive antigenic diversity of *Ot* strains and an incomplete understanding of protective immunity^13–15^.

IFN-γ is a key immunoregulatory cytokine for host protection against intracellular pathogens^16,17^. Upon binding to its receptor, IFN-γ activates the Janus kinase-signal transducer and activator of transcription 1 signaling pathway, leading to the induction of interferon-stimulated genes (ISGs) that suppress pathogen replication and enhance the expression of immune effector and signaling proteins. *Ot* infection elicits robust IFN-γ responses in scrub typhus patients and murine models^18–25^. It is reported that scrub typhus patients exhibit markedly upregulated IFN-γ and Th1-associated cytokine CXCL10^18^. Using established inbred C57BL/6 and outbred CD-1 mouse models, our group has consistently demonstrated that *Ot* infection elicits robust IFN-γ responses, accompanied by the induction of CXCL10, across lymphoid and parenchymal organs^22,26^. More recently, we used a panel of transgenic mouse models and demonstrated a critical role of IFN-γ, but not IFN-Ι, in host protection against *Ot* infection^27^. We found that *Ifngr1*-deficient mice were highly susceptible to infection and developed eschar-like lesions, a hallmark feature observed in a proportion of scrub typhus patients^27–29^. Although we have demonstrated a critical role of IFN-γ in scrub typhus, the mechanism underlying IFN-γ signaling-mediated control of *Ot* infection remains unclear.

Inducible nitric oxide synthase (iNOS, encoded by *Nos2*), typically induced following STAT1 activation, is a well-characterized antimicrobial effector molecule^30^. IFN-γ-STAT1 signaling directly and indirectly promotes iNOS transcription through cooperative interactions with additional transcription factors, including interferon regulatory factor-1 and NF-κB^31^. This results in the production of nitric oxide (NO) that contributes to pathogen clearance by exerting antimicrobial and immunomodulatory effects, particularly in activated M1 macrophages^32,33^. *Ot*, formerly called *Rickettsia tsutsugamushi*, is closely related to the *Rickettsia* genus; both infect endothelial and immune cells, leading to systemic inflammation, vascular dysfunction, and potentially severe multi-organ disease^34^. In *Rickettsia rickettsii* infection, NO-producing macrophages restricted bacterial growth *in vitro*, whereas pharmacological inhibition of iNOS abrogated this antimicrobial effect^35^. Similar findings have been reported in *Rickettsia prowazekii* infection, in which NO was shown to prevent bacterial invasion of host cells through direct bactericidal activity^36^. During *Rickettsia parkeri* infection, IFN-γ induces *Nos2* expression in mouse primary macrophages, leading to the inhibition of bacterial growth *in vitro*^37^. In *Ot* infection, iNOS-expressing macrophages have been observed within infiltrative parenchymal nodules of multiple organs in a scrub typhus mouse model^38^. Consistently, *in vitro* studies demonstrated that IFN-γ inhibits *Ot* growth in mouse bone marrow-derived macrophages (BMDMs), whereas pharmacological inhibition of iNOS reverses the antibacterial effects of IFN-γ^38^. Our previous study also demonstrated that *Ot* infection can significantly increase *Nos2* expression in mouse primary macrophages *in vitro* at 4 and 24 hours post-infection as compared with the mock control^24^. Although these studies highlight the antimicrobial role of iNOS *in vitro*, its contribution to bacterial control and host immune homeostasis *in vivo* remains controversial. For example, both protective and nonessential roles for iNOS have been reported in mouse models of *Mycobacterium tuberculosis* infection^39,40^, highlighting the context-dependent and complex functions of iNOS *in vivo*. Given our demonstration of the critical role of IFN-γ in controlling *Ot* infection *in vivo*^27,41^, we sought to define the contribution of iNOS to *Ot* control and disease outcome in the mouse model of scrub typhus.

In this study, we employed our established intradermal scrub typhus mouse model in combination with transgenic mouse strains. Our results demonstrate that iNOS is required for restricting *Ot* growth in murine macrophages *in vitro*. Notably, *Nos2*^−/−^ mice developed eschars like those observed in *Ifngr1*^−/−^ mice, indicating a critical role for the IFN-γ/iNOS axis in preventing skin lesion formation. Although *Nos2*^−/−^ mice showed comparable bacterial burdens as compared to wild-type (WT) mice during acute infection, they exhibited higher inflammatory responses in organs and dysregulated systemic homeostasis relative to WT controls. More importantly, the absence of *Nos2* resulted in delayed bacterial clearance during the recovery phase, leading to persistent inflammatory responses in various organs. Collectively, these findings define an essential role for iNOS in *Ot* infection and provide new insight into host protective immunity in scrub typhus.

## Materials and Methods

### Animals, infection, and treatment

C57BL/6j (#000664), *Ifngr1*^−/−^ (#003288), and *Nos2*^−/−^ (#002609) mice were purchased from Jackson Laboratory. All tissue processing and analysis procedures were performed in the BSL3 or BSL2 facilities. All procedures were approved by the Institutional Animal Care and Use Committee (IACUC #2101001A) and the Institutional Biosafety Committee, in accordance with Guidelines for Biosafety in Microbiological and Biomedical Laboratories. UTMB operates to comply with the United States Department of Agriculture (USDA) Animal Welfare Act (Public Law 89-544), the Health Research Extension Act of 1985 (Public Law 99-158), the Public Health Service Policy on Humane Care and Use of Laboratory Animals, and the NAS Guide for the Care and Use of Laboratory Animals (ISBN-13). UTMB is a registered Research Facility under the Animal Welfare Act and has a current assurance on file with the Office of Laboratory Animal Welfare, in compliance with NIH Policy.

For animal infection, mice were first anesthetized in a chamber connected to a VetFlo isoflurane vaporizer. Next, mice were carefully removed from the chamber, and the infection site (right flank) was shaved using an electric trimmer. Mice were inoculated with *Ot* Karp strain (2×10^3^ FFU, 20 µL volume) in the dermis of the flank by using a 0.3-mL insulin syringe with 31-G needles (Sol-Millennium, Chicago, IL), and were monitored daily for weight loss, skin lesion, signs of disease, and survival. At indicated time-points, mice were euthanized by CO_2_ inhalation, followed by cardiac perfusion with 10 mL of PBS. Tissues/blood were harvested for further analysis. Eschar-like lesions were photographed, and lesion size was measured using a vernier caliper in the ABSL3 biosafety cabinet.

### Bacterial Stock Preparation

Bacteria were inoculated onto L929 murine fibroblast monolayers cultured in T175 flasks and incubated at 37°C with gentle rocking for 2 h. After incubation, Minimum Essential Medium supplemented with 10% fetal bovine serum, 100 U/mL penicillin, and 100 μg/mL streptomycin was added. At 7 days post-infection (dpi), cells were harvested by scraping, resuspended in MEM, and lysed by vortexing with 0.5-mm glass beads (MP Biomedicals, Solon, OH) for 1 min. The lysate was centrifuged at 300 × g for 10 min to pellet cellular debris and glass beads. The supernatant was transferred to Oak Ridge high-speed centrifuge bottles and centrifuged at 22,000 × g for 45 min at 4°C to pellet bacteria. The bacterial pellet was resuspended in sucrose phosphate glutamate buffer (0.218 M sucrose, 3.8 mM KH₂PO₄, 7.2 mM K₂HPO₄, 4.9 mM monosodium L-glutamate, pH 7.0) to prepare bacterial stocks, which were stored at −80°C. The same batch of bacterial stocks was used for all experiments described in this study. Bacterial titers were determined by focus-forming assay as previously described^22,42^.

### Quantitative PCR (qPCR) for Measuring Bacterial Burdens

Genomic DNA from tissues and cultured cells was extracted using the DNeasy Blood and Tissue Kit (Qiagen, Germantown, MD) and subjected to qPCR analysis, as previously described^22,42^. The 47-kDa gene was amplified using the primer pair OtsuF630 (5 ′ - AACTGATTTTATTCAAACTAATGCTGCT-3 ′) and OtsuR747 (5 ′ - TATGCCTGAGTAAGATACGTGAATGGAATT-3 ′) (IDT, Coralville, IA). Detection was performed with the probe OtsuPr665 (5 ′ - 6FAM-TGGGTAGCTTTGGTGGACCGATGTTTAATCT-TAMRA-3 ′) (IDT) using SsoAdvanced Universal Probes Supermix (Bio-Rad, Hercules, CA). Bacterial loads were normalized to the total DNA concentration or each well for tissues or cultured cells, respectively. Absolute quantification was determined using a 10-fold serial dilution of an *Ot* Karp 47-kDa plasmid standard, which contains a single copy of the target gene.

### Quantitative Reverse Transcriptase PCR (qRT-PCR)

Total RNA was extracted from mouse tissues using the RNeasy Mini Kit (Qiagen). RNA concentration and purity were assessed using a BioTek microplate reader; cDNA was synthesized from 1 μg of total RNA using the iScript Reverse Transcription Kit (Bio-Rad). qRT-PCR reactions (10 μL in total volume) contained 5 μL of iTaq SYBR Green Supermix (Bio-Rad), 1 μL of forward and reverse primer mix (final concentration 0.5 μM each), and 4 μL of diluted cDNA. Amplification was carried out on a CFX96 Touch Real-Time PCR Detection System (Bio-Rad) with the following cycling conditions: initial denaturation at 95°C for 30s, followed by 40 cycles of 95°C for 15s and 60°C for 60s. Relative gene expression levels were calculated using the 2^−ΔΔCt^ method and normalized to glyceraldehyde-3-phosphate dehydrogenase (*Gapdh*) as the internal control. All primers were synthesized by Integrated DNA Technologies (IDT, Coralville, IA) and are listed in **Supplementary Table 1**. Primer sequences were obtained from PrimerBank.

### Histology

Tissues were fixed in 10% neutral buffered formalin and paraffin-embedded at the UTMB Research Histology Service Core. Paraffin sections (5-μm thick) were prepared and stained with hematoxylin and eosin (H&E). Images were acquired using an Olympus BX53 microscope, and at least five randomly selected fields per section were captured.

### Clinical Chemistry Parameters

Animal blood chemistry analysis was performed by using the VetScan Chemistry Analyzer (Zoetis, Parsippany-Troy Hills, NJ). Briefly, mouse serum was separated from the blood using the BD MicroTainer Blood Collection Tube with SST Amber and was uploaded into the VetScan Comprehensive Diagnostic Profile reagent rotor (100 μL/rotor). This rotor is used for quantitative determinations of alanine aminotransferase (ALT), albumin (ALB), alkaline phosphatase (ALP), amylase (AMY), globulin (GLOB), glucose (GLU), potassium (K^+^), sodium (Na^+^) and total protein (TP). This analysis was performed in the UTMB ABSL3 animal facility, as in our previous reports^27,41^.

### Bone Marrow-Derived Macrophage Generation and Infection

BMDMs were generated from WT B6 and *Nos2*^−/−^ mice by culturing bone marrow cells with recombinant M-CSF (40 ng/ml, Biolegend, San Diago, CA), as in our previous reports^24,43^. To polarize M1 macrophages, BMDMs were treated with lipopolysaccharide (LPS, 100 ng/mL, Sigma, St. Louis, MO) and IFN-γ (20 ng/mL, Biolegend, San Diago, CA) for 16 h. Cells were infected with *Ot* Karp at multiplicities of infection (MOI 2 and 10) for 2 h, followed by thorough washing with warm PBS and culturing in fresh medium. In some experiments, serial concentrations (10, 100 or 1000 µM) of the selective iNOS inhibitor 1400W were used during M1 polarization and infection to inhibit iNOS signaling. Cells were collected at 24, 48 and 72 h post-infection for bacterial burden measurement by qPCR.

### Immunofluorescence

BMDMs were seeded on sterile glass coverslips placed in 24-well plates, followed by M1 polarization and *Ot* infection, as described above. At 72 h post-infection, cells were washed with PBS, fixed with 4% paraformaldehyde (15 min at room temperature), and permeabilized with 0.1% Triton X-100 in PBS for 10 min. Non-specific binding sites were blocked by incubating the samples in blocking buffer containing 5% bovine serum albumin in PBS for 1 h at room temperature. Following blocking, cells were incubated overnight at 4°C with mouse anti-*Ot* TSA47 monoclonal antibody (Yao-Hong Biotechnology, Taiwan, 1:2000 dilution). Goat anti-mouse IgG-AF488 (1: 2000 dilution, Southern Biotech, Birmingham, AL) were then applied and incubated for 1 h at room temperature in the dark. Cell nuclei were counterstained with 4’,6-diamidino-2-phenylindole (DAPI) for 5 min. At least five images were captured per coverslip using an Olympus BX53 fluorescence microscope.

### Flow Cytometry

Spleen single-cell suspensions were prepared directly by pressing spleen tissues through 70-μm cell strainers using a sterile 3-cc syringe. Equivalent portions of lung tissues were harvested from mice, minced, and digested with 0.05% collagenase type IV (Thermo Fisher Scientific, Waltham, MA) in RPMI-1640 medium for 30 min at 37°C. Minced tissues were loaded into Medicons and homogenized using a BD Mediamachine System (BD Biosciences, Franklin Lakes, NJ). Lung single-cell suspensions were made by passing lung homogenates through 70-μm cell strainers. Red blood cells were removed by using Red Cell Lysis Buffer (Sigma-Aldrich, St. Louis, MO). For surface marker analysis, leukocytes were stained with the Fixable Viability Dye (eFluor 506, Thermo Fisher Scientific) for live/dead cell discrimination, blocked with FcγR blocker, and incubated with fluorochrome-labeled antibodies for 30 min at 4°C. The fluorochrome-labeled antibodies were purchased from Thermo Fisher Scientific or Biolegend (San Diego, CA) as below: Alexa Fluor 700 anti-CD11b (M1/70), APC anti-Ly6G (1A8), PE/Dazzle-594 anti-Ly6C (HK1.4), Percp-Cy5.5 anti-CD11c (N418), FITC anti-MHCII (M5/114.15.2), PE-Cy7 anti-CD3ε (145-2C11), Percp-Cy5.5 anti-CD4 (GK1.5), APC-Cy7-anti-CD8a (53-6.7), BV711 anti-CD44 (IM7), APC anti-CD62L (MEL-14), PE CF594 anti-NK1.1 (PK136), FITC anti-CD69 (H1.2F3), PE-Cy7 anti-CD80 (16-10A1), PE anti-CD103 (2E7), BV785 anti-B220 (RA3-6B2), Alexa Fluor 488 anti-GL7 (GL7), Percp-Cy5.5 anti-CD38 (90), BV510 anti-IgD (11-26c.2a), BV421 anti-PDL2 (TY25) and BV605 anti-CD73 (TY/11.8). BV421 anti-F4/80 (T45-2342) and RB780 anti-CD3ε (500A2) were purchased from BD Bioscience. For iNOS staining, BMDMs were blocked with FcγR blocker and fixed by using IC buffer (Thermo Fisher Scientific) for 15 min. After fixation, cells were washed by permeabilization buffer and stained with PE-Cy7 anti-iNOS (CXNFT, Thermo Fisher Scientific) for 45 min at 4°C. All cell samples were fixed in 2% paraformaldehyde overnight at 4°C, acquired by a BD Symphony A5 and analyzed via FlowJo software version 10 (BD Bioscience). The gating strategy of immune cell subsets was based on recent publications^27,41,44,45^.

### Statistical analysis

Data are presented as mean ± standard deviation (SD). One-way ANOVA was used for multiple group comparisons, while unpaired t-test was used for two group comparisons. Two-way ANOVA was used for bodyweight change analysis. All data were analyzed by using GraphPad Prism software 10. Statistically significant values are denoted as *, *p* < 0.05; **, *p* < 0.01; ***, *p* < 0.001 and ****, *p* < 0.0001; ns, not significant, respectively.

## Results

### Dynamic patterns of *Ifng*/*Stat1*/*Nos2* expression in mouse tissues following *Ot* infection

We have previously reported that *Ot*-infected mice develop a type 1-skewed immune response, as evidenced by markedly elevated IFN-γ expression^22,23,46^. To characterize the temporal dynamics of iNOS-related signaling during infection, we intradermally infected mice with the *Ot* Karp strain and quantified *Ifng*, *Stat1*, and *Nos2* transcript levels in multiple organs at indicated time points (**Figure 1**). qRT-PCR analysis revealed significant upregulation of *Ifng* expression in the spleen, draining lymph nodes (dLN), lungs, and liver on 7 or 14 dpi. While *Stat1* levels in the spleen and dLN were modestly increased, their levels in the lungs and liver were markedly upregulated. *Nos2* expression was also significantly upregulated across multiple tissues, with particularly strong and early induction in the liver. We also found that *Nos2* upregulation in the lungs was relatively delayed and was primarily evident at 14 dpi. Notably, elevated *Nos2* expression persisted in both the liver and spleen at 42 dpi, indicating that iNOS may contribute to host defense during the acute phase, as well as to immune regulation during the recovery phase of *Ot* infection.

**Figure 1.**
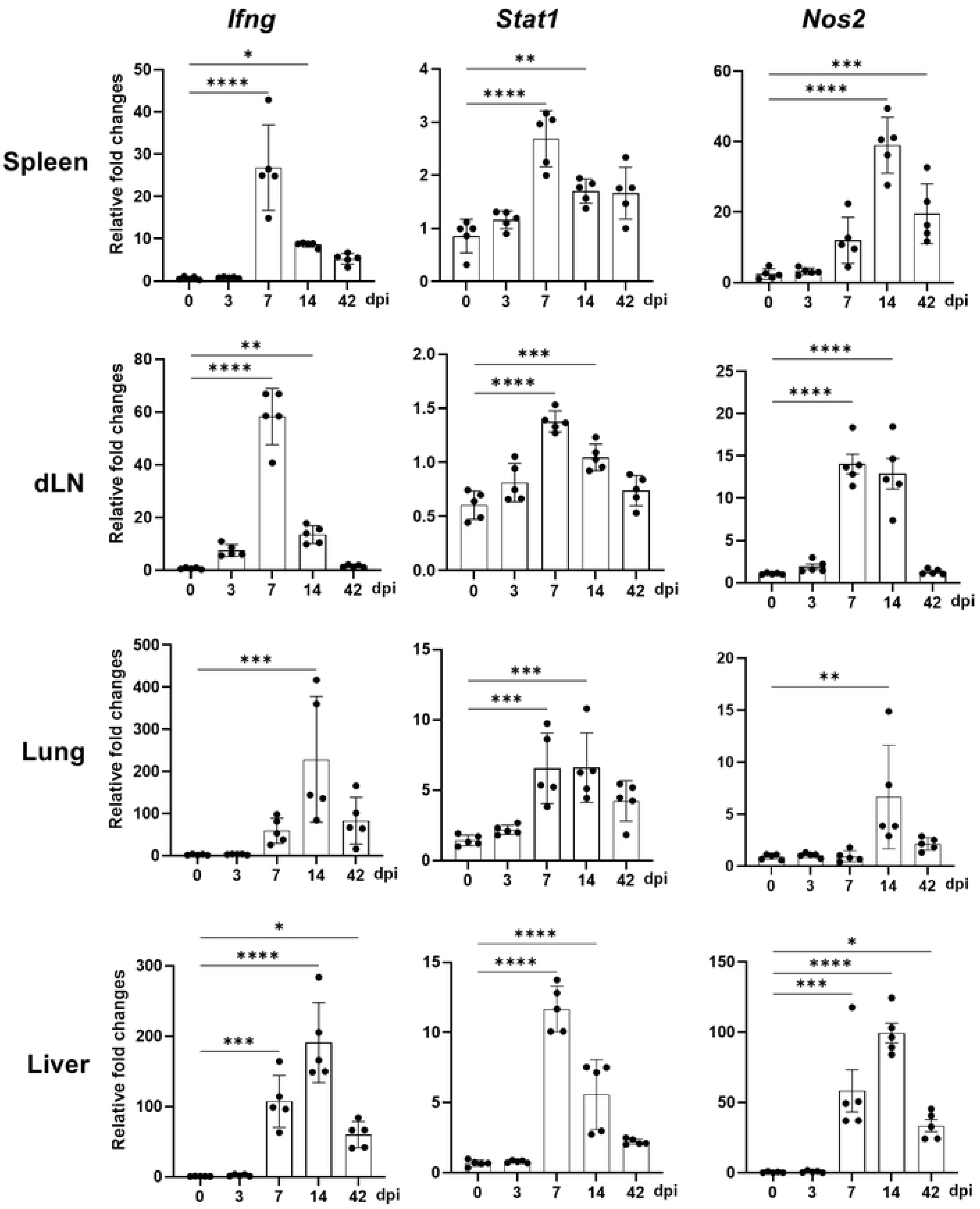
*Ot* infection induced *Nos2* and its related gene expression in mouse organs. B6 mice (n = 5/group) were intradermally infected with *Ot* (2 × 10^3^ FFU) on the flank. Mouse tissues were harvested on 3, 7, 14 and 42 days post-infection (dpi). Mice receiving PBS were used as control (day 0). RNA was extracted from spleen, dLN, lungs and liver, followed by cDNA synthesis. The transcript levels of IFN-γ, Stat1 and Nos2 were analyzed by qRT-PCR. The values are shown as mean ± SD from single experiments and are representative of two independent experiments. One-way ANOVA followed by Dunnett’s multiple-comparisons test was used to compare infected samples with the control group. *, *p* < 0.05; **, *p* < 0.01; ***, *p* < 0.001; ****, *p* < 0.0001. Data without any statistical difference were not labeled.

### iNOS signaling contributes to the restriction of *Ot* growth in mouse macrophages

To assess the functional role of iNOS in macrophage-mediated control of *Ot*, we generated BMDMs from WT and *Nos2*^−/−^ mice and stimulated them with IFN-γ alone or in combination with LPS for 24 h to induce M1 polarization. Flow cytometric analysis revealed that IFN-γ alone induced only limited iNOS expression in WT macrophages, whereas combined stimulation with IFN-γ and LPS resulted in robust iNOS induction (**Supplementary Fig. 1**). We next infected these cells with *Ot* at MOIs of 2 and 10, and quantified bacterial burdens at 1-3 dpi. As shown in **Figure 2A**, WT macrophages stimulated with IFN-γ and LPS exhibited a significantly enhanced capacity to restrict bacterial growth at both MOIs compared with untreated controls, as evidenced by reduced bacterial burdens at all time points. Importantly, the enhanced antibacterial activity observed in activated macrophages was abolished in the absence of iNOS, as *Nos2*^−/−^ macrophages exhibited significantly higher bacterial burdens compared with WT controls. To further evaluate the contribution of iNOS to *Ot* growth control *in vitro*, M1 macrophages were treated with increasing concentrations of the selective iNOS inhibitor 1400W. No overt cytotoxicity was observed in M1 macrophages at inhibitor concentrations up to 1 mM (**Supplementary Fig. 2**). We then infected these cells at MOI 2 and 10 and measured bacterial burdens at 3 dpi. As expected, M1 macrophages markedly inhibited *Ot* growth compared with M0 macrophages (**Figure 2B**). Notably, pharmacological inhibition of iNOS partially reversed this restriction, resulting in higher bacterial burdens compared with untreated M1 macrophages, though still lower than those observed in M0 cells (**Figure 2B**). To further confirm our findings, we infected BMDMs with *Ot* at an MOI of 2 and performed IF staining of bacteria at 3 dpi. As shown in **Figure 2C**, M0 macrophages exhibited significantly stronger fluorescent bacterial staining compared to M1 macrophages after *Ot* infection, indicating that M1 macrophages could effectively inhibit bacterial growth. Notably, blocking iNOS impaired the suppressive effects of M1 macrophages in controlling *Ot* growth. These results demonstrate that iNOS contributes to the anti-*Ot* activity in mouse M1 macrophages.

**Figure 2.**
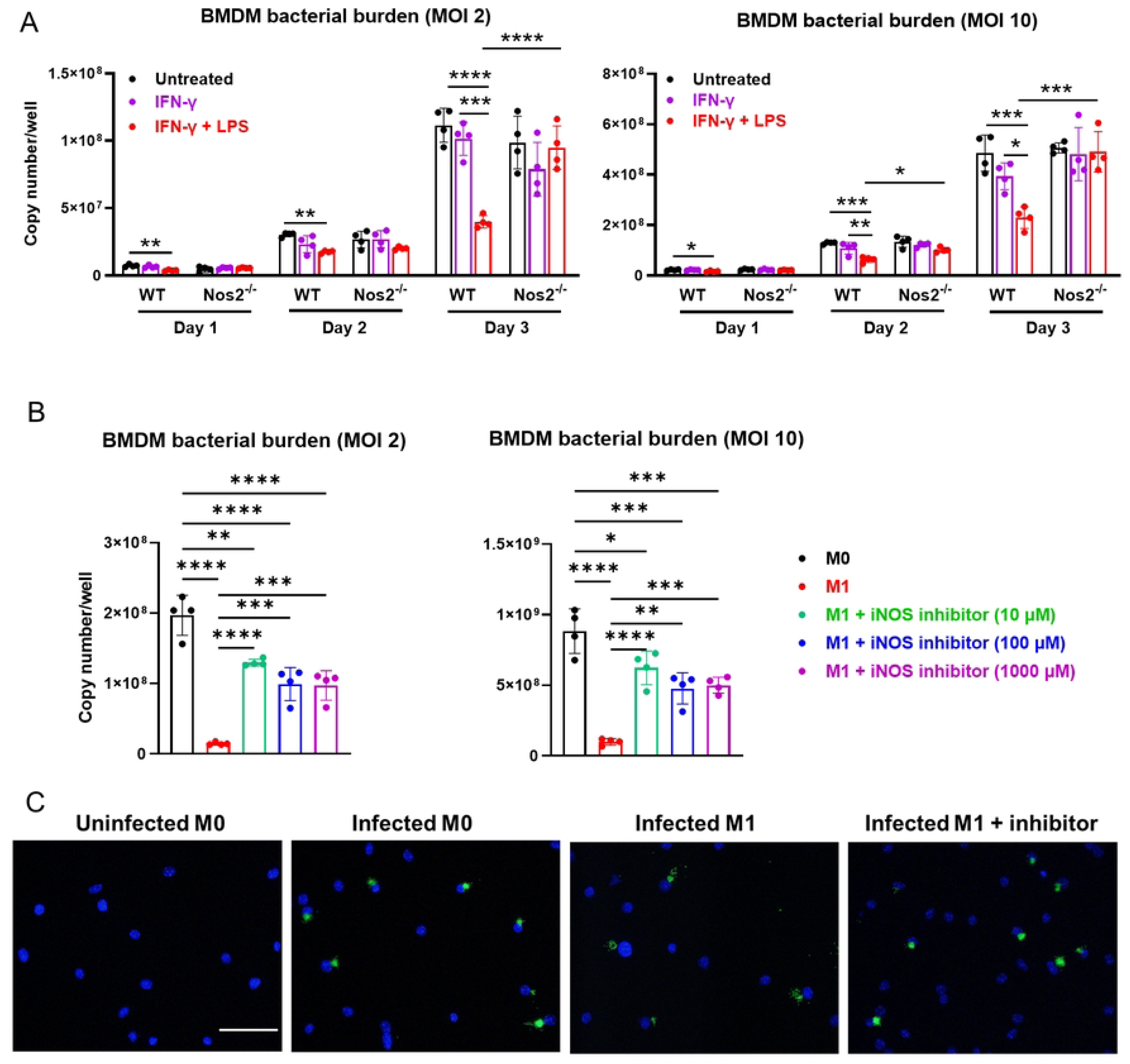
iNOS signaling was required for M1 macrophage-mediated restriction of *Ot* growth *in vitro*. (A) Bone marrow cells were isolated from WT or *Nos2*^−/−^ mice, followed by the differentiation into macrophages *in vitro*. Macrophages were polarized with either IFN-γ (20 ng/mL) alone or IFN-γ plus LPS (100 ng/mL) for 16 h, followed by *Ot* infection (MOI 2 and 10). Cells were harvested on day 1, 2 and 3 post-infection for bacterial burden measurement using qPCR. (B) M1 macrophages were treated with indicated doses of iNOS inhibitor 1400W, followed by *Ot* infection (MOI 2 and 10). M0 macrophages were used as a control. Bacterial burdens on day 3 post-infection were quantified by qPCR. The values are shown as mean ± SD from single experiments and are representative of two independent experiments. One-way ANOVA followed by Tukey’s multiple comparisons test was used to compare each group. *, *p* < 0.05; **, *p* < 0.01; ***, *p* < 0.001; ****, *p* < 0.0001. Data without any statistical difference were not labeled. (C) M0 and M1 macrophages were infected with *Ot* (MOI 10) and were cultured in the presence or absence of the iNOS inhibitor 1400W (10 µM). Uninfected M0 macrophages were used as a control. Cells were fixed on day 3 post-infection and stained with anti-*Ot* TSA47 antibody (green color) and DAPI (blue color). The representative images of two independent experiments were shown.

### *Nos2^−/−^* mice showed increased inflammatory responses and the formation of eschar-like lesions

We next determined the role of iNOS *in vivo* during *Ot* infection. WT and *Nos2*^−/−^ mice were intradermally infected with 2,000 FFU of the *Ot* Karp strain. *Ifngr1*^−/−^ mice were included as a susceptible control, as in our previous report^27^. We found that *Nos2*^−/−^ mice exhibited slightly more weight loss compared to WT mice over the 6-week observation period (**Figure 3A**). No mortality was observed in either WT or *Nos2*^−/−^ mice (**Figure 3B**), suggesting that both strains were resistant to intradermal *Ot* infection. As expected, *Ifngr1*^−/−^ mice began losing weight at 10 dpi, and all succumbed to infection by 17 dpi (**Figure 3A and 3B**). Notably, *Nos2*^−/−^ mice, but not WT mice, formed obvious eschar-like lesions at 14 dpi with lesion sizes comparable to those observed in *Ifngr1*^−/−^ mice (**Figure 3C**). This finding suggests that the IFN-γ/iNOS signaling axis might be a key determinant of eschar formation. Furthermore, the dLN of WT mice were enlarged during the acute phase of infection (7 and 14 dpi) but contracted at the recovery phase (42 dpi), when bacterial burden was largely resolved. In contrast, *Nos2*^−/−^ mice maintained enlarged dLN at 42 dpi (**Figure 3D**), indicating persistent inflammation in the absence of *Nos2* in *Ot* infection.

**Figure 3.**
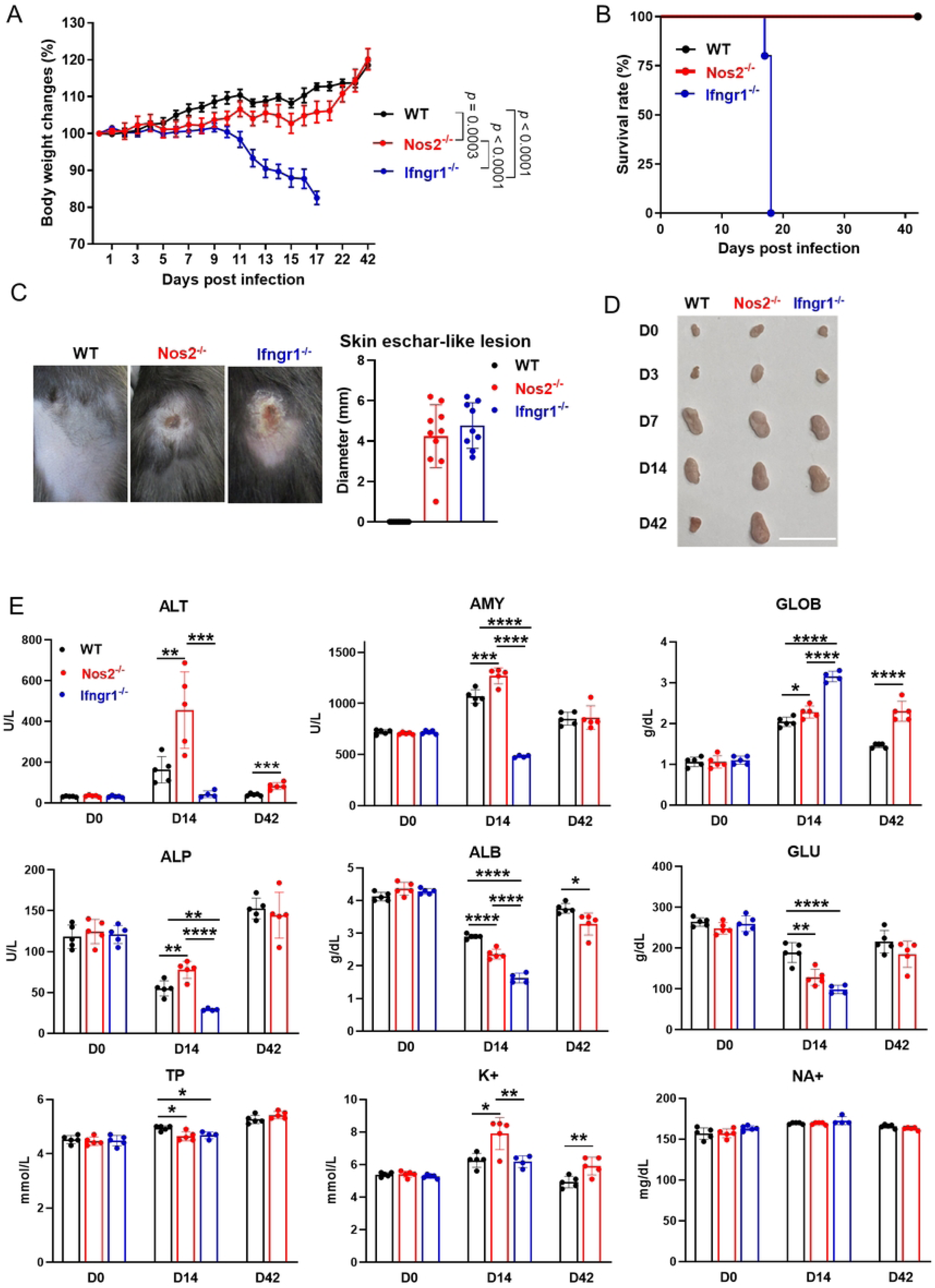
Deficiency of Nos2 was dispensable for disease progression, but resulted in eschar-like lesions in *Ot*-infected mice. WT, *Nos2*^−/−^ and *Ifngr1*^−/−^ mice (n = 5/group) were infected as described in Figure 1 and were monitored daily for (A) bodyweight changes and (B) survival rates. (C) Photographs of eschar-like lesions were taken at 13 dpi. The size of the lesion was measured by using a vernier scale. (D) The photos of dLN were taken at different time points. (E) Serum chemistry parameters were detected by using VetScan Comprehensive Diagnostic Profile Reagent Rotor. Data are shown as mean ± SD from two independent experiments. The data of bodyweight changes were analyzed by two-way ANOVA with a Tukey’s multiple comparisons test. The skin lesion sizes and serum chemistry parameters were analyzed by one-way ANOVA with a Tukey’s multiple comparisons test for three groups (day 0 and 14), while unpaired t-test was used for comparison of two groups (day 42). *, *p* < 0.05; **, *p* < 0.01; ***, *p* < 0.001; ****, *p* < 0.0001. Data without any statistical difference were not labeled.

To comprehensively evaluate the physiological status of mice during infection, we measured the dynamic changes in serum biochemical parameters using VetScan Comprehensive Diagnostic Profile reagent rotors (**Figure 3E**). At 14 dpi, *Nos2*^−/−^ mice exhibited significantly higher levels of ALT, AMY, GLOB, and ALP compared to WT mice. These findings suggest that the absence of *Nos2* exacerbates organ inflammation, particularly liver injury, during acute *Ot* infection. Similar to *Ifngr1*^−/−^ mice, *Nos2*^−/−^ mice displayed reduced levels of ALB, GLU, and TP, indicating dysregulated physiological and metabolic homeostasis in these deficient mice. Moreover, *Nos2*^−/−^ mice continued to show elevated ALT and GLOB levels at 42 dpi, alongside decreased ALB compared to WT controls. This lower ratio of ALB-to-GLOB in *Nos2*^−/−^ mice may suggest persistent inflammation during the recovery phase^47–49^.

Because *Nos2*^−/−^ mice did not exhibit significant disease manifestations compared to WT controls, we sought to determine whether bacterial clearance differed across stages of infection. We quantified *Ot* burdens by qPCR (**Figure 4A**) and found that *Nos2*^−/−^ mice displayed comparable bacterial loads in the lungs, spleen, and dLN at 3, 7, and 14 dpi. As expected, *Ifngr1*^−/−^ mice failed to control *Ot* infection in these organs. Notably, at 42 dpi, bacterial burdens remained detectable in all three organs of *Nos2*^−/−^ mice, whereas they were almost undetectable in WT mice (**Figure 4A**). We further performed histological assessment of lung and liver tissues. As shown in **Figure 4B and C**, *Ot* infection resulted in similar levels of hepatic inflammatory infiltration at 7 and 14 dpi in both WT and *Nos2*^−/−^ mice, whereas *Ifngr1*^−/−^ mice exhibited severe liver damage as evidenced by significant hepatic necrosis at 14 dpi. In addition, the perivascular infiltrations in the lungs were comparable between WT and *Nos2*^−/−^ mice during the acute infection. Notably, inflammatory infiltration was markedly resolved in WT mice at 42 dpi, whereas *Nos2*^−/−^ mice continued to exhibit persistent inflammation in the affected organs. These results suggest that *Nos2* may play a central role in bacterial clearance and resolution of inflammation during the host recovery stage.

**Figure 4.**
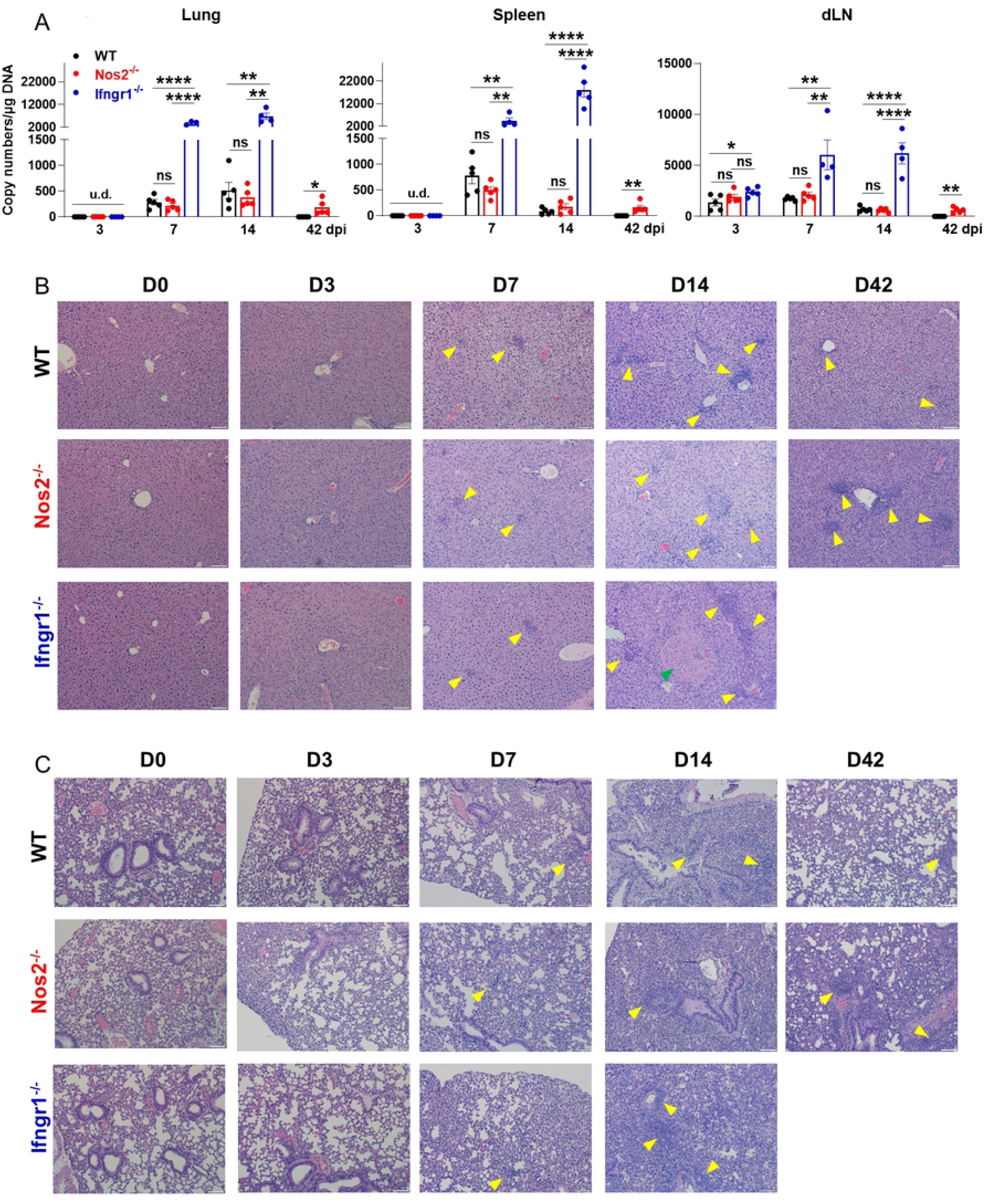
*Nos2*^−/−^ mice exhibited increased bacterial burden and exacerbated tissue pathology compared with WT mice at the recovery stage. **WT**, *Nos2*^−/−^ and *Ifngr1*^−/−^ mice (n = 5/group) were infected as described in Figure 1. (A) Lungs, spleen and dLN were collected at indicated timepoints for bacterial burden measurement by qPCR. Data are shown as mean ± SD from two independent experiments and were analyzed by one-way ANOVA with a Tukey’s multiple comparisons test at indicated timepoints. *, *p* < 0.05; **, *p* < 0.01; ****, *p* < 0.0001. ns, not significant. Representative histological images of (B) livers and (C) lungs were shown. Yellow arrows indicate inflammatory infiltration. Green arrow indicates liver necrosis. Scale bar = 100 µm.

### *Nos2* deficiency results in increased activation of neutrophils and macrophages in acute infection

To determine the role of iNOS in regulating immune responses during *Ot* infection, we analyzed the dynamic profile of splenic immune cell activation by flow cytometry (**Supplementary Figure 3)**. As shown in **Figure 5A-D**, *Nos2*^−/−^ mice exhibited comparable numbers of total splenocytes, activated T cells, and activated NK cells compared to WT mice during the acute phase of infection. In contrast, *Ifngr1*^−/−^ mice displayed increased numbers of activated T cells but reduced activated NK cells, likely reflecting their substantially higher bacterial burden and severely impaired IFN signaling, respectively. Consistent with *Ifngr1* deficiency, *Nos2* deficiency led to a reduction in Ly6C^hi^ monocytes but a marked increase in activated neutrophils (**Figure 5E-F**). Notably, at 14 dpi, the absolute number of neutrophils was much higher than Ly6C^hi^ monocytes in all mouse strains, suggesting that neutrophils may represent a dominant inflammatory population contributing to dysregulated physiological and metabolic homeostasis. Furthermore, *Nos2* deficiency did not significantly alter the total numbers of dendritic cells or macrophages in the spleen (**Figure 5G-H**). However, unlike *Ifngr1*^−/−^ mice, which exhibited impaired myeloid cell activation, *Nos2*^−/−^ mice showed comparable or even elevated myeloid activation, as evidenced by increased CD80 expression relative to WT controls. We also isolated cells from lungs and analyzed the dynamic profile of pulmonary immune activation (**Supplementary Figure 4**). Consistent with the finding in the spleen, there were no major changes in T cell and NK cell activation between WT and *Nos2*^−/−^ mice at 0, 3, 7 and 14 dpi (**Supplementary Figure 4A-C**). However, *Nos2* deficiency resulted in elevated lung macrophage activation at 14 dpi as evidenced by increased CD80 and MHCII levels on macrophages of *Nos2*^−/−^ mice (**Supplementary Figure 4D**). Therefore, these results suggest that *Nos2* deficiency may lead to increased myeloid cell activation and upregulated inflammatory responses in the acute infection of *Ot*.

**Figure 5.**
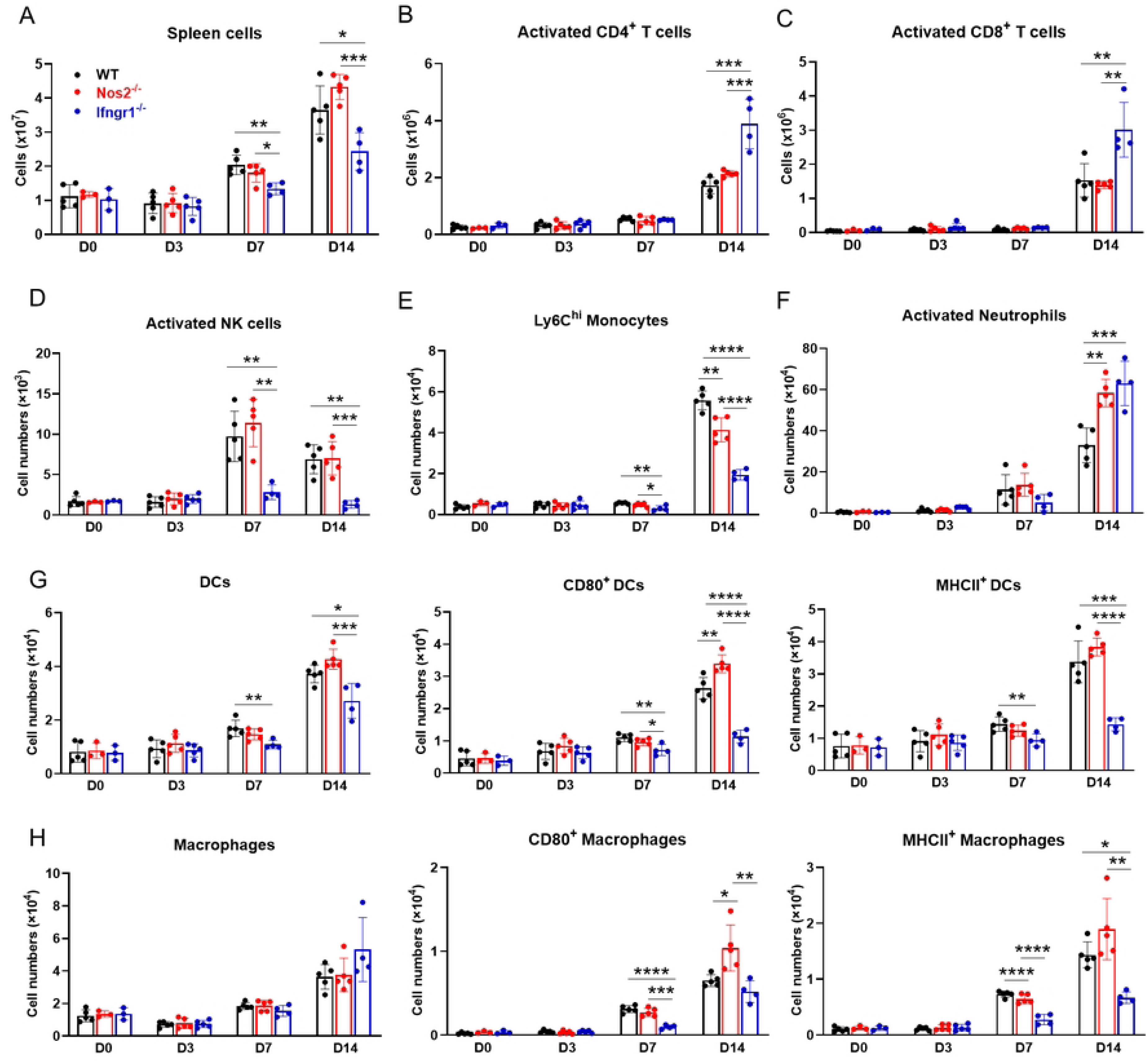
*Nos2*^−/−^ mice displayed altered myeloid cell response compared with WT mice during the acute stage of infection. WT, *Nos2*^−/−^ and *Ifngr1*^−/−^ mice (n = 5/group) were infected as described in Figure 1. Single cell suspensions were prepared from the spleens at days 0, 3, 7 and 14, followed by the flow cytometric analysis. Absolute numbers of immune cell subsets were presented. (A) Total spleen cells. (B) CD44^+^CD62L^−^ activated CD4^+^ T cells. (C) CD44^+^CD62L^−^activated CD8^+^ T cells. (D) CD69^+^ activated NK cells. (E) CD11b^+^Ly6C^hi^ monocytes. (F) CD63^+^ activated neutrophils. (G) CD11c^+^ dendritic cells; CD80^+^ dendritic cells; MHCII^+^ dendritic cells. (H) F4/80^+^ macrophages; CD80^+^ macrophages; MHCII^+^ macrophages. Data are shown as mean ± SD from two independent experiments and were analyzed by one-way ANOVA with a Tukey’s multiple comparisons test at indicated timepoints. *, *p* < 0.05; **, *p* < 0.01; ***, *p* < 0.001; ****, *p* < 0.0001. Data without any statistical difference were not labeled.

### *Nos2* deficiency leads to persistent inflammation in the recovery phase of *Ot* infection

Given enlarged lymphoid organs, sustained bacterial burdens, and uncontracted tissue inflammation in *Nos2*^−/−^ mice at 42 dpi (**Figure 3D, 4A-C**), we sought to evaluate the host immune homeostasis at this recovery stage. We analyzed the WT and *Nos2*^−/−^ mouse spleen samples by flow cytometry. No samples from *Ifngr1*^−/−^ mice are available, as all of them succumbed to infection at 17 dpi. Our flow cytometric results showed that the percentages of activated CD4^+^ and CD8^+^ T cells were two-fold higher in *Nos2*^−/−^ mice compared to WT mice (**Figure 6A**). Consistently, the absolute numbers of activated CD4⁺ and CD8⁺ T cells were increased by more than threefold in *Nos2*^−/−^ mice. In addition, *Nos2* deficiency caused approximately a three-fold increase in the total number of neutrophils and CD63⁺ activated neutrophils (**Figure 6B**). Moreover, macrophage and Ly6C^hi^ monocyte numbers were also significantly elevated by approximately three-fold in *Nos2*^−/−^ mice (**Figure 6C and D**). More importantly, the expression of CD80 was significantly higher on both *Nos2*^−/−^ macrophages and Ly6C^hi^ monocytes, indicating the upregulated myeloid cell activation in the absence of *Nos2*. We also assessed B cell populations in the spleen, as in our previous report^44^. Although B cell percentages were comparable between the two groups, *Nos2*^−/−^ mice had significantly higher B cell numbers, likely due to their significant increase of total spleen cells (**Supplementary Figure 5**). Furthermore, *Nos2* deficiency resulted in increased B cell activation as evidenced by increased germinal center B cells and memory B cells (**Supplementary Figure 5**). In addition to the spleen, we also analyzed immune responses in lungs, a major organ of *Ot* replication and pathogenesis. Our lung results were consistent with the spleen data, showing significantly increased T cell activation, as evidenced by the higher percentages of CD44^+^CD62^−^subpopulations (**Supplementary Figure 6A**). Interestingly, *Nos2*^−/−^ mice exhibited reduced tissue-resident memory CD8^+^ T cells, as demonstrated by decreased percentages of CD69^+^CD103^+^ subset (**Supplementary Figure 6B**). Although the percentages of lung macrophages and Ly6C^hi^ monocytes were comparable between WT and *Nos2*^−/−^ mice, the expression of CD80 and MHCII on macrophages was significantly higher in *Nos2*^−/−^ mice (**Supplementary Figure 6C)**. Unlike in the spleens, activated neutrophils in the lungs were comparable between WT and *Nos2*^−/−^ mice (**Supplementary Figure 6D)**, which may suggest distinct immune dynamics across different organs.

**Fig. 6.**
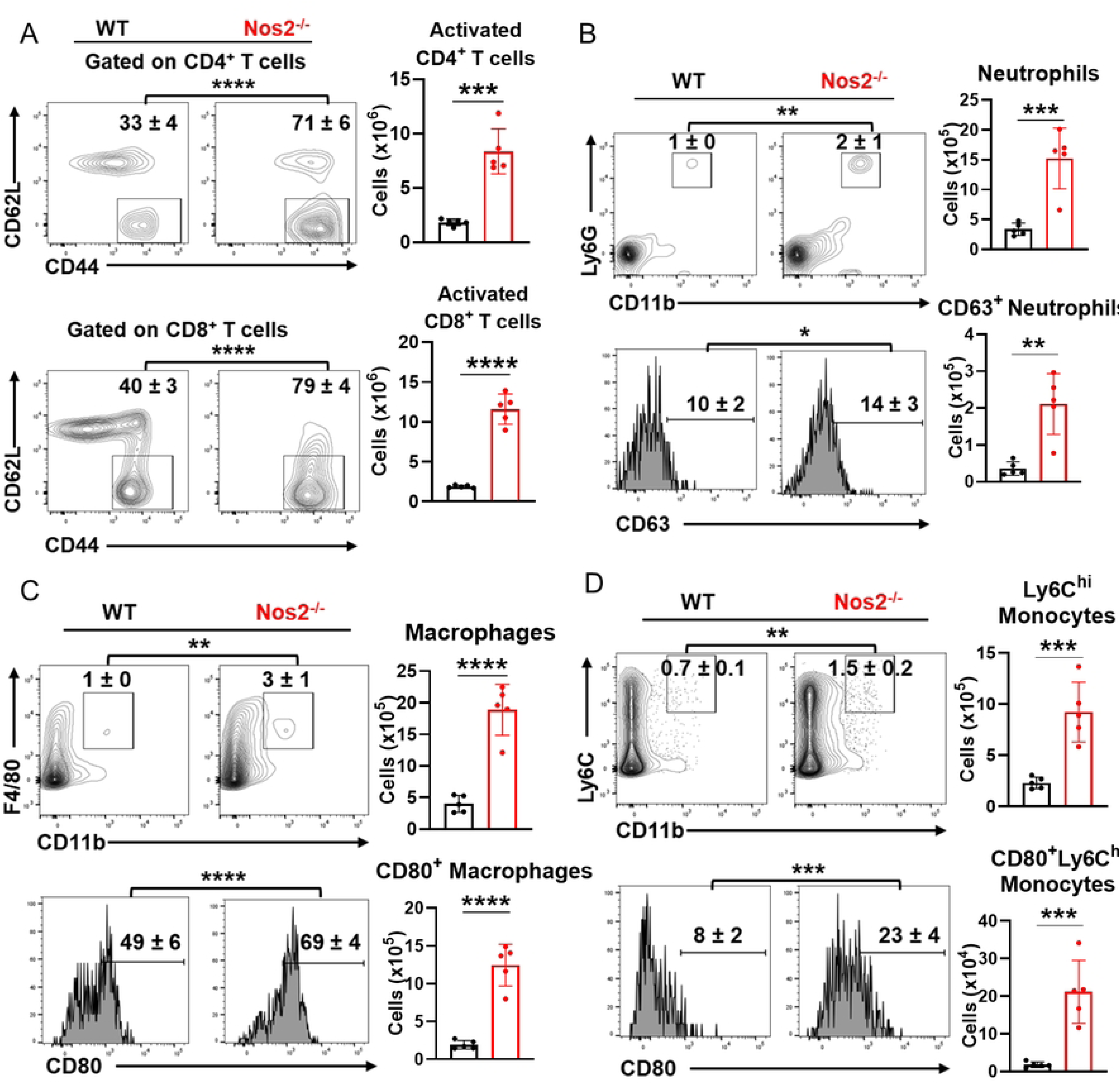
N*o*s2 deficiency resulted in persistent immune activation at the recovery phase of *Ot* infection. WT, *Nos2*^−/−^ and *Ifngr1*^−/−^ mice (n = 5/group) were infected as described in Figure 1. Single cell suspensions were prepared from the spleens on day 42, followed by the flow cytometric analysis. All *Ifngr1*^−/−^ mice succumbed to infection before day 42. Both cell percentages and absolute numbers of immune cell subsets were shown. (A) CD44^+^CD62L^−^ activated CD4^+^ and CD8^+^ T cells. (B) Total neutrophils and CD63^+^ activated neutrophils. (C) Total macrophages and CD80^+^ macrophages. (D) Total monocytes and CD80^+^ monocytes. Data are shown as mean ± SD from two independent experiments and were analyzed by unpaired t-test. *, *p* < 0.05; **, *p* < 0.01; ***, *p* < 0.001; ****, *p* < 0.0001. Data without any statistical difference were not labeled.

To further support our findings, we used qRT-PCR to examine key inflammatory gene profiles in dLN, liver, spleen, and lungs during the infection. As shown in **Figure 7A**, *Ot* infection upregulated several inflammatory genes in dLN, including *Il1b*, *Il6*, *Il27*, *Ccl2*, *Cxcl9*, and *Cxcl10*. During the acute phase of infection, *Nos2* deficiency was associated with elevated *Cxcl10* at 3 dpi, *Il1b* and *Il6* at 7 dpi, and *Ccl2* at 14 dpi. Strikingly, at 42 dpi, *Nos2*^−/−^ mice exhibited upregulation of all inflammatory genes examined, except *Tnf* in dLN, whereas WT mice showed resolution and contraction of inflammatory responses. Consistently, at 42 dpi, the expression levels of all these inflammatory genes were significantly elevated in the livers of *Nos2*^−/−^ mice (**Figure 7B**). This sustained hepatic inflammation at the recovery stage may contribute to the dysregulated physiological homeostasis reflected in the altered serum biochemical parameters (**Figure 3E**). In addition, several inflammatory genes (*Ccl2*, *Tnf*, and *Cxcl10,* etc.) were also upregulated in the spleen and/or lung tissues at 42 dpi (**Supplementary Figure 7)**. In all, these findings indicate systemic and persistent inflammatory responses in *Nos2*^−/−^ mice during the recovery phase of *Ot* infection.

**Figure 7.**
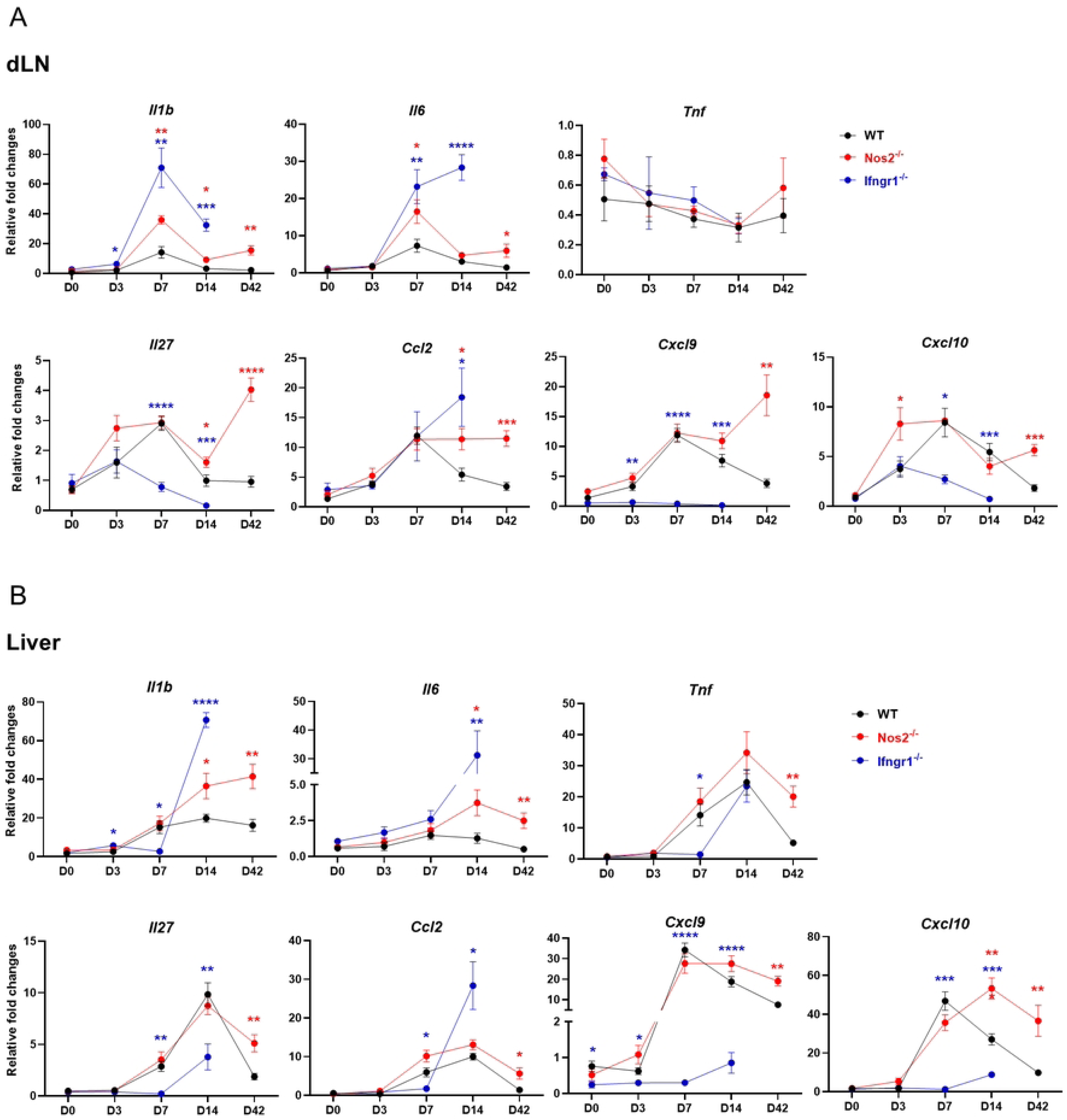
*Nos2* deficiency resulted in unresolved inflammation in mouse organs at the recovery phase of *Ot* infection. WT, *Nos2*^−/−^ and *Ifngr1*^−/−^ mice (n = 5/group) were infected as described in Figure 1. RNA was extracted from various organs and the dynamic patterns of inflammatory genes, including *Il1b*, *Il6*, *Tnf*, *Il27*, *Ccl2*, *Cxcl9* and *Cxcl10* in (A) dLN and (B) livers were assessed by qRT-PCR. Data are shown as mean ± SD from two independent experiments and were analyzed by one-way ANOVA with a Tukey’s multiple comparisons test at indicated timepoints. *, *p* < 0.05; **, *p* < 0.01; ***, *p* < 0.001; ****, *p* < 0.0001. The red stats represent comparison between *Nos2*^−/−^ and WT, while the blue stars represent comparison between *Ifngr1*^−/−^ and WT. Data without any statistical difference were not labeled.

## Discussion

iNOS, which is expressed in response to proinflammatory stimuli (e.g., LPS, IFN-γ and TNF-α), acts as a crucial mediator of immune activation, inflammation, and host defense^50^. As a hallmark functional marker of M1 macrophages, iNOS is primarily induced through the classical IFN-γ/STAT1 pathway in synergy with pattern-recognition receptor signaling (e.g., TLRs)^51^. In infectious diseases, the role of iNOS has been investigated extensively, suggesting that iNOS can act as a double-edged sword^52–54^. While iNOS-derived NO mediates pathogen killing and exerts anti-inflammatory effects^30,33,35,55,56^, iNOS can also drive excessive inflammation, contributing to blood-brain barrier disruption in bacterial meningitis^57^, increased mortality and morbidity during influenza A infection^58^, and heightened susceptibility to fungal infections^59^.

The antibacterial role of iNOS has been well documented in studies of *Rickettsia* species, which are closely related to *Ot*. Activated, nitric oxide-producing macrophages restrict *Rickettsia rickettsii* replication, whereas pharmacologic inhibition of iNOS abolishes this growth restriction^35^. Similarly, early *Rickettsia prowazekii* infection is suppressed in L929 cells treated with IFN-γ plus TNF-α, and this effect is reversed by iNOS inhibition^36^. More recently, IFN-I has been shown to induce iNOS expression in an IRF5-dependent manner, contributing to control of *Rickettsia parkeri* infection^60^. In sharp contrast, very little is known regarding the precise role of iNOS against *Ot*, even though *Ot* infection is known to elicit type 1-skewed immune responses in human patients and in different mouse models^18,22,46^. Elevated IFN-γ, in combination with proinflammatory cytokines such as TNF-α, can potently induce iNOS expression^61^. We previously demonstrated *in vivo* that TNF-α and IFN-γ signals are critical for host protection against *Ot* infection^27,45^, indicating downstream molecules such as iNOS may play a role in controlling *Ot* infection. In scrub typhus patients, iNOS expression is markedly increased, particularly in alveolar macrophages and damaged lung tissues^62^. Consistently, histochemical analyses reveal iNOS expression across multiple organs, where it colocalizes with activated macrophages/microglia in murine models of scrub typhus^38^. Collectively, these reports from human studies and experimental animal models support a critical role for iNOS in mediating host protective responses against *Ot* infection.

In this study, we aimed to define the functional role of iNOS in *Ot* infection using our established intradermal infection model. We first characterized the kinetics of iNOS expression across organs during infection. qRT-PCR analysis revealed parallel expression patterns of *Ifng*, *Stat1*, and *Nos2* in multiple tissues (Figure 1). Consistent with the previous report^38^, *Nos2* expression peaked around 14 dpi, with the highest levels observed in the liver. Notably, *Nos2* expression remained elevated in the spleen and liver at 42 dpi, suggesting a potential role for iNOS in the recovery and resolution phase of infection. We next employed two complementary approaches (*Nos2*^−/−^ cells and iNOS selective inhibitor treatment) to assess the anti-*Ot* function of iNOS in macrophages (Figure 2). Our results demonstrated that iNOS is required for effective restriction of bacterial growth in activated macrophages. In line with these findings, immunostaining using an anti-TSA47 monoclonal antibody showed reduced bacterial burden in M1 macrophages compared to M0 controls, whereas inhibition of iNOS restored bacterial replication in M1 macrophages. Collectively, these *in vitro* results support and extend prior observations that iNOS is a critical effector mediating antimicrobial activity against *Ot* in activated macrophages.

Next, we infected WT and *Nos2^−/−^* mice via intradermal inoculation with *Ot* and monitored the disease outcomes. Although *Nos2^−/−^* mice exhibited greater weight loss compared to WT controls, both strains survived infection with only mild disease symptoms, whereas all *Ifngr1^−/−^* mice succumbed around 16-17 dpi (Figure 3). Interestingly, despite their resistance to mortality, *Nos2^−/−^* mice developed eschar-like lesions of comparable size to those observed in *Ifngr1^−/−^* mice (Figure 3C). These findings suggest that iNOS is not essential for animal survival during *Ot* infection but plays a critical role in regulating skin pathology. Although skin lesions appeared reduced during the recovery stage, accurate evaluation was difficult due to hair regrowth at the inoculation site and the potential for skin damage if the area was re-shaved. Interestingly, a recent study also reported eschar lesions in *Nos2^−/−^* mice infected with *Rickettsia parkeri*^37^. Consistently, iNOS has been implicated in wound healing and tissue repair in other skin injury models, including chemical-induced, excisional and diabetic wounds^63–66^. Our data therefore support an IFN-γ/iNOS axis as a key determinant of skin eschar formation in scrub typhus. Notably, we also observed similar skin lesions in our newly established humanized IFN-γ mouse model, but not in WT mice^41^. Given that IFN-γ and iNOS signaling are generally less robust in humans than in mice^37,41^, these observations suggest that eschar formation in scrub typhus patients may result, at least in part, from relatively limited IFN-γ/iNOS activity in the skin following chigger bite.

Persistence of *Ot* has been reported in scrub typhus patients^67,68^; however, the mechanisms by which *Ot* evades host immune defenses to establish persistence remain unclear. Although we observed comparable bacterial burdens and pathology between WT and *Nos2*^−/−^ mice in multiple organs during the acute phase (3, 7, and 14 dpi), *Nos2*^−/−^ mice showed significantly enlarged lymphoid organs at 42 dpi, suggesting delayed immune contraction in the absence of iNOS signaling (Figure 3D). More importantly, bacterial burdens remained detectable in the lungs, spleen, and dLN of *Nos2*^−/−^ mice, but were undetectable in WT controls at this late stage (Figure 4A). This result indicates that while iNOS contributes only modestly to *Ot* control during the acute stage, it is critical for complete pathogen clearance and immune contraction at the recovery stage. Flow cytometric analysis further revealed persistent immune activation in the spleen and lungs of *Nos2^−/−^* mice at 42 dpi, characterized by increased activation and infiltration of T cells, macrophages, neutrophils, and monocytes (Figure 6). Consistently, transcriptional profiling showed elevated expression of proinflammatory genes, including *Il1b*, *Il6*, *Il27*, *Ccl2*, *Cxcl9*, and *Cxcl10*, in *Nos2^−/−^* mice compared to WT controls at the recovery stage, indicating sustained systemic inflammation in the absence of iNOS signals (Figure 7). In addition, histopathological analysis corroborated these findings, demonstrating increased inflammatory infiltration and tissue damage in the lungs and liver of *Nos2^−/−^* mice (Figure 4B-C). This was further supported by serum chemistry, which showed elevated ALT levels and reduced ALB and GLU in *Nos2^−/−^* mice relative to WT mice (Figure 3E). Collectively, these data suggest that the absence of iNOS signals may lead to persistent bacterial infection and prolonged proinflammatory responses during the recovery stage of *Ot* infection.

Although we found that iNOS was required for activated macrophage-mediated restriction of *Ot in vitro* (Figure 2), its apparently dispensable role in restricting *Ot* during the acute phase *in vivo* remains unclear. Based on our findings and recent studies, we propose several explanations. First, *Ot* may evade macrophage-mediated innate defense *in vivo*. Following intradermal infection, although macrophages can phagocytose bacteria, dendritic cells appear to be the primary carriers that transport bacteria from the skin to dLN^69^. This trafficking route may allow the pathogen to evade macrophage-mediated killing, effectively infect lymphatic endothelial cells, and ultimately disseminate into visceral organs. Second, optimal iNOS induction typically requires both pathogen-derived signals (e.g., LPS) and proinflammatory cytokines such as IFN-γ, TNF-α, and IL-1β. However, *Ot* lacks LPS^70^, a potent inducer of iNOS and NO production. As a result, NO levels in the skin and lymphatic tissues may be insufficient during early infection, enabling bacterial growth and immune evasion. Third, T cell-mediated immunity, particularly CD8⁺ T cells, plays a dominant role in controlling infection *in vivo*^71,72^. Although iNOS deficiency has been demonstrated to enhance Th1 responses during bacterial and viral infection^73–75^, we did not find major differences in T cell or NK cell activation between WT and *Nos2*^−/−^ mice during the acute phase (Figure 5), suggesting that iNOS may play a marginal role in shaping early lymphocyte responses in this context. Finally, *Ot* infection induces robust type 1 immune responses and upregulates numerous ISGs. Certain ISGs, such as members of the guanylate-binding protein family^41^, may serve as key antimicrobial effectors that compensate for the loss of iNOS signals^60^. Further studies are needed to identify the critical ISGs responsible for bacterial control and to define their underlying mechanisms.

Our study has several limitations that warrant further studies. The intradermal inoculation used here mimics the natural route of *Ot* infection, but it is a non-lethal model in WT B6 mouse models^38,76,77^. It remains unclear as to how iNOS-mediated responses contribute to *Ot* clearance and/or cellular injury in lethal models (e.g., intravenous inoculation)^22,42^. Given the known differences in human and murine cells in their capacity to induce iNOS^37,78^, it will be important to examine the role of iNOS in the control of *Ot* growth in human cells (e.g., macrophages). Since *Ot* can replicate in endothelial cells, which produce relatively low NO concentrations via eNOS for vascular function and metabolic homeostasis^79,80^, it will be interesting to examine whether iNOS-derived NO can help control *Ot* infection in endothelial cells. Future studies are still needed to elucidate the mechanisms by which iNOS contributes to *Ot* clearance and disease outcomes. Collectively, our findings reveal the first *in vivo* role of an IFN-γ/iNOS axis in scrub typhus and offer new insights into host immune defense against *Ot* infection.

## Acknowledgments

We would like to thank the UTMB Flow Cytometry & Cell Sorting Core Lab, as well as Dr. David Walker and Nicole Mendell for generously sharing reagents and facilities.

---

**Supplementary Figure 1. iNOS expression in WT and *Nos2*^−/−^ BMDMs were analyzed by flow cytometry.** Bone marrow cells were isolated from WT or *Nos2*^−/−^ mice, followed by the differentiation into macrophages *in vitro*. Macrophages were polarized with either IFN-γ (20 ng/mL) alone or IFN-γ plus LPS (100 ng/mL) for 16 h. Intracellular flow cytometry was performed to measure iNOS expression. The percentages of iNOS^+^ cells were shown on the flow panels. Data are shown as mean ± SD from two independent experiments.

**Supplementary Figure 2. iNOS inhibitor 1400W did not show any obvious cytotoxicity in BMDMs.** BMDMs were cultured with serial concentrations (10, 100 or 1000 µM) of the iNOS inhibitor 1400W for 3 days. The images were captured by EVOS M5000 Imaging System. Scale bar: 50 µm. Results are from two independent experiments.

**Supplementary Figure 3. Gating strategy of flow cytometry.** Single cells were first gated based on FSC-A and SSC-A, followed by exclusion of doublets using FSC-A vs. FSC-H. Dead cells were excluded using a fixable Live/Dead viability dye. NK cells were identified as CD3⁻NK1.1⁺ cells, and CD69 was used as a marker of NK cell activation. Activated T cells were defined based on CD44 and CD62L expression, while CD69⁺CD103⁺ T cells were identified as lung resident memory T cell subsets. Neutrophils were gated as CD11b⁺Ly6G⁺ cells, and CD63 was used as a marker of neutrophil activation. Dendritic cells, monocytes, and macrophages were identified based on CD11c⁺, Ly6C^hi^, and F4/80⁺ expression, respectively. Germinal center (GC) B cells were identified as B220⁺CD38⁻GL-7⁺ cells. For memory B cell analysis, non-GC B cells were first selected, followed by gating on IgD⁻CD73⁺ cells. These IgD⁻CD73⁺ cells were further gated for CD80⁺PD-L2⁺ expression to define the memory B cell subset.

**Supplementary Figure 4. Flow cytometric percentages of splenic immune cells during the acute stage of infection.** WT, *Nos2*^−/−^ and *Ifngr1*^−/−^ mice (n = 5/group) were infected as described in Figure 1. Single cell suspensions were prepared from the spleens at days 0, 3, 7 and 14, followed by the flow cytometric analysis. Percentages of immune cell subsets were presented. (A) NK cells and CD69^+^ activated NK cells. (B) CD4^+^ T cells and CD44^+^CD62L^−^ activated CD4^+^ T cells. (C) CD8^+^ T cells and CD44^+^CD62L^−^ activated CD8^+^ T cells. (D) CD80^+^ macrophages and MHCII^+^ macrophages. (E) Neutrophils and CD63^+^ activated neutrophils. (F) CD80^+^ dendritic cells and MHCII^+^ dendritic cells. Data are shown as mean ± SD from two independent experiments and were analyzed by one-way ANOVA with a Tukey’s multiple comparisons test at indicated timepoints. *, *p* < 0.05; **, *p* < 0.01; ***, *p* < 0.001; ****, *p* < 0.0001. Data without any statistical difference were not labeled.

**Supplementary Figure 5. *Nos2*^−/−^ mice exhibited elevated B cell responses during the recovery stage.** WT and *Nos2*^−/−^ mice (n = 5/group) were infected as described in Figure 1. Single cell suspensions were prepared from the spleens on day 42, followed by the flow cytometric analysis. Both cell percentages and absolute numbers of (A) B cells, (B) GC B cells, and (C) memory B cells were shown. Data are shown as mean ± SD from two independent experiments and were analyzed by unpaired t-test. **, *p* < 0.01; ***, *p* < 0.001. Data without any statistical difference were not labeled.

Supplementary Figure 6. *Nos2* deficiency resulted in altered immune homeostasis in the lung at the recovery phase of *Ot* infection. WT and *Nos2*^−/−^ mice (n = 5/group) were infected as described in Figure 1. Single cell suspensions were prepared from lungs on day 42, followed by the flow cytometric analysis. (A) Activated CD4^+^ and CD8^+^ T cells. (B) Lung resident memory T cells. (C) Monocytes and macrophages were gated on Ly6c^hi^ and F4/80. The expression levels of CD80 and MHCII on macrophages were analyzed. (D) Neutrophils and CD63^+^ activated neutrophils. Data are shown as mean ± SD from two independent experiments and were analyzed by unpaired t-test. **, *p* < 0.01; ***, *p* < 0.001; ****, *p* < 0.0001. ns, not significant.

**Supplementary Figure 7. Dynamic profile of inflammatory genes in mouse spleen and lungs at the recovery phase of *Ot* infection.** WT, *Nos2*^−/−^ and *Ifngr1*^−/−^ mice (n = 5/group) were infected as described in Figure 1. RNA was extracted from (A) spleen and (B) lungs and the dynamic patterns of inflammatory genes, including *Il1b*, *Il6*, *Tnf*, *Il27*, *Ccl2*, *Cxcl9* and *Cxcl10* were assessed by qRT-PCR. Data are shown as mean ± SD from two independent experiments and were analyzed by one-way ANOVA with a Tukey’s multiple comparisons test at indicated timepoints. *, *p* < 0.05; **, *p* < 0.01; ***, *p* < 0.001; ****, *p* < 0.0001. The red stats represent comparison between *Nos2*^−/−^ and WT, while the blue stars represent comparison between *Ifngr1*^−/−^ and WT. Data without any statistical difference were not labeled.

## References

1. Lee CS, Hwang JH, Lee HB, Kwon KS. Risk factors leading to fatal outcome in scrub typhus patients. Am J Trop Med Hyg. 2009;81(3):484–488.

2. Paris DH, Shelite TR, Day NP, Walker DH. Unresolved problems related to scrub typhus: a seriously neglected life-threatening disease. Am J Trop Med Hyg. 2013;89(2):301–307.

3. Tilak R, Kunte R. Scrub typhus strikes back: Are we ready? Med J Armed Forces India. 2019;75(1):8–17.

4. Jiang J, Richards AL. Scrub Typhus: No Longer Restricted to the Tsutsugamushi Triangle. Trop Med Infect Dis. 2018;3(1):11.

5. Xu G, Walker DH, Jupiter D, Melby PC, Arcari CM. A review of the global epidemiology of scrub typhus. PLoS Negl Trop Dis. 2017;11(11):e0006062.

6. Wang Q, Ma T, Ding F, et al. Global and regional seroprevalence, incidence, mortality of, and risk factors for scrub typhus: A systematic review and meta-analysis. Int J Infect Dis. 2024;146:107151.

7. Chen K, Travanty NV, Garshong R, et al. Detection of *Orientia* spp. Bacteria in Field-Collected Free-Living *Eutrombicula* Chigger Mites, United States. Emerg Infect Dis. 2023;29(8):1676–1679.

8. Abernathy HA, Ursery L, Merdjane BA, Giandomenico DA, Boyce RM. *Orientia tsutsugamushi* Antibodies in Patients with Eschars and Suspected Tickborne Disease. Emerg Infect Dis. 2025;31(11):2187–2190.

9. Peter JV, Sudarsan TI, Prakash JA, Varghese GM. Severe scrub typhus infection: Clinical features, diagnostic challenges and management. World J Crit Care Med. 2015;4(3):244–250.

10. Long J, He Y, Li K, Wu X, Zhang Z, Wei Y. Antibiotic therapy failure and clinical outcomes in scrub typhus patients from Guangzhou city, southern China. Sci Rep. 2026;16(1):7555.

11. Prakash JAJ. Scrub typhus: risks, diagnostic issues, and management challenges. Res Rep Trop Med. 2017;8:73–83.

12. Song X, Xie S, Huang X, Chen Z. The diagnosis and treatment of scrub typhus should be emphasized in non-endemic areas: A retrospective case series study. Medicine (Baltimore). 2023;102(8):e32988.

13. Kelly DJ, Fuerst PA, Ching WM, Richards AL. Scrub typhus: the geographic distribution of phenotypic and genotypic variants of *Orientia tsutsugamushi*. Clin Infect Dis. 2009;48 Suppl 3:S203–230.

14. Walker DH, Mendell NL. A scrub typhus vaccine presents a challenging unmet need. NPJ Vaccines. 2023;8(1):11.

15. Chaichana P, Satapoomin N, Kullapanich C, et al. Comparative virulence analysis of seven diverse strains of *Orientia tsutsugamushi* reveals a multifaceted and complex interplay of virulence factors responsible for disease. PLoS Pathog. 2025;21(6):e1012833.

16. Casanova JL, MacMicking JD, Nathan CF. Interferon-gamma and infectious diseases: Lessons and prospects. Science. 2024;384(6693):eadl2016.

17. Kak G, Raza M, Tiwari BK. Interferon-gamma (IFN-gamma): Exploring its implications in infectious diseases. Biomol Concepts. 2018;9(1):64–79.

18. Munch CC, Upadhaya BP, Rayamajhee B, et al. Multiple *Orientia* clusters and Th1-skewed chemokine profile: a cross-sectional study in patients with scrub typhus from Nepal. Int J Infect Dis. 2023;128:78–87.

19. Ge H, Farris CM, Tong M, Maina A, Richards AL. Transcriptional profiles of cytokines and chemokines reveal important pro-inflammatory response from endothelial cells during *Orientia tsutsugamushi* infection. Microbes Infect. 2019;21(7):313–320.

20. Moron CG, Popov VL, Feng HM, Wear D, Walker DH. Identification of the target cells of *Orientia tsutsugamushi* in human cases of scrub typhus. Mod Pathol. 2001;14(8):752–759.

21. Chierakul W, de Fost M, Suputtamongkol Y, et al. Differential expression of interferon-gamma and interferon-gamma-inducing cytokines in Thai patients with scrub typhus or leptospirosis. Clin Immunol. 2004;113(2):140–144.

22. Soong L, Wang H, Shelite TR, et al. Strong type 1, but impaired type 2, immune responses contribute to *Orientia tsutsugamushi*-induced pathology in mice. PLoS Negl Trop Dis. 2014;8(9):e3191.

23. Soong L, Shelite TR, Xing Y, et al. Type 1-skewed neuroinflammation and vascular damage associated with *Orientia tsutsugamushi* infection in mice. PLoS Negl Trop Dis. 2017;11(7):e0005765.

24. Fisher J, Card G, Liang Y, Trent B, Rosenzweig H, Soong L. *Orientia tsutsugamushi* selectively stimulates the C-type lectin receptor Mincle and type 1-skewed proinflammatory immune responses. PLoS Pathog. 2021;17(7):e1009782.

25. Inthawong M, Sunyakumthorn P, Wongwairot S, et al. A time-course comparative clinical and immune response evaluation study between the human pathogenic *Orientia tsutsugamushi* strains: Karp and Gilliam in a rhesus macaque (Macaca mulatta) model. PLoS Negl Trop Dis. 2022;16(8):e0010611.

26. Thiriot J, Liang Y, Fisher J, Walker DH, Soong L. Host transcriptomic profiling of CD-1 outbred mice with severe clinical outcomes following infection with *Orientia tsutsugamushi*. PLoS Negl Trop Dis. 2022;16(11):e0010459.

27. Liang Y, Wang H, Sun K, Sun J, Soong L. Lack of the IFN-gamma signal leads to lethal *Orientia tsutsugamushi* infection in mice with skin eschar lesions. PLoS Pathog. 2024;20(5):e1012020.

28. Kundavaram AP, Jonathan AJ, Nathaniel SD, Varghese GM. Eschar in scrub typhus: a valuable clue to the diagnosis. J Postgrad Med. 2013;59(3):177–178.

29. Park J, Woo SH, Lee CS. Evolution of Eschar in Scrub Typhus. Am J Trop Med Hyg. 2016;95(6):1223–1224.

30. Chakravortty D, Hensel M. Inducible nitric oxide synthase and control of intracellular bacterial pathogens. Microbes Infect. 2003;5(7):621–627.

31. Farlik M, Reutterer B, Schindler C, et al. Nonconventional initiation complex assembly by STAT and NF-kappaB transcription factors regulates nitric oxide synthase expression. Immunity. 2010;33(1):25–34.

32. Okda M, Spina S, Safaee Fakhr B, Carroll RW. The antimicrobial effects of nitric oxide: A narrative review. Nitric Oxide. 2025;155:20–40.

33. Nathan CF, Hibbs JB, Jr. Role of nitric oxide synthesis in macrophage antimicrobial activity. Curr Opin Immunol. 1991;3(1):65–70.

34. Richards AL, Jiang J. Scrub Typhus: Historic Perspective and Current Status of the Worldwide Presence of *Orientia* Species. Trop Med Infect Dis. 2020;5(2):49.

35. Fitzsimmons LF, Clark TR, Hackstadt T. Nitric Oxide Inhibition of *Rickettsia rickettsii*. Infect Immun. 2021;89(12):e0037121.

36. Turco J, Liu H, Gottlieb SF, Winkler HH. Nitric oxide-mediated inhibition of the ability of *Rickettsia prowazekii* to infect mouse fibroblasts and mouse macrophagelike cells. Infect Immun. 1998;66(2):558–566.

37. Luu AP, Guzman AA, Bouin A, Drayman N, Burke TP. Differences between human and rodent nitric oxide production dictate susceptibility to tick-borne *Rickettsia*. bioRxiv. 2025:2025.2006.2027.661835.

38. Keller CA, Hauptmann M, Kolbaum J, et al. Dissemination of *Orientia tsutsugamushi* and inflammatory responses in a murine model of scrub typhus. PLoS Negl Trop Dis. 2014;8(8):e3064.

39. MacMicking JD, North RJ, LaCourse R, Mudgett JS, Shah SK, Nathan CF. Identification of nitric oxide synthase as a protective locus against tuberculosis. Proc Natl Acad Sci U S A. 1997;94(10):5243–5248.

40. Cooper AM, Pearl JE, Brooks JV, Ehlers S, Orme IM. Expression of the nitric oxide synthase 2 gene is not essential for early control of *Mycobacterium tuberculosis* in the murine lung. Infect Immun. 2000;68(12):6879–6882.

41. Cho RH, Gao L, Wang H, et al. A humanized IFN-gamma mouse model reveals skin eschar formation, enhanced susceptibility and scrub typhus pathogenesis. PLoS Pathog. 2026;22(2):e1013419.

42. Shelite TR, Saito TB, Mendell NL, et al. Hematogenously disseminated *Orientia tsutsugamushi*-infected murine model of scrub typhus [corrected]. PLoS Negl Trop Dis. 2014;8(7):e2966.

43. Trent B, Liang Y, Xing Y, et al. Polarized lung inflammation and Tie2/angiopoietin-mediated endothelial dysfunction during severe *Orientia tsutsugamushi* infection. PLoS Negl Trop Dis. 2020;14(3):e0007675.

44. Gonzales C, Liang Y, Thiriot J, et al. Unique B Cell and Germinal Center Responses in Mice with Severe versus Mild *Orientia tsutsugamushi* Infection. bioRxiv. 2025:2025.2003.2027.645674.

45. Liang Y, Fisher J, Gonzales C, et al. Distinct Role of TNFR1 and TNFR2 in Protective Immunity Against *Orientia tsutsugamushi* Infection in Mice. Front Immunol. 2022;13:867924.

46. Soong L. Dysregulated Th1 Immune and Vascular Responses in Scrub Typhus Pathogenesis. J Immunol. 2018;200(4):1233–1240.

47. Gabay C, Kushner I. Acute-phase proteins and other systemic responses to inflammation. N Engl J Med. 1999;340(6):448–454.

48. Park J, Kim HJ, Kim J, Choi YB, Shin YS, Lee MJ. Predictive value of serum albumin-to-globulin ratio for incident chronic kidney disease: A 12-year community-based prospective study. PLoS One. 2020;15(9):e0238421.

49. Ulloque-Badaracco JR, Mosquera-Rojas MD, Hernandez-Bustamante EA, Alarcon-Braga EA, Herrera-Anazco P, Benites-Zapata VA. Prognostic value of albumin-to-globulin ratio in COVID-19 patients: A systematic review and meta-analysis. Heliyon. 2022;8(5):e09457.

50. Forstermann U, Sessa WC. Nitric oxide synthases: regulation and function. Eur Heart J. 2012;33(7):829–837.

51. Kang JH, Kim JH, Gim JA, Lee MY. iNOS in Macrophage Polarization: Pharmacological and Regulatory Insights. Int J Mol Sci. 2025;26(24):12056.

52. Masood M, Singh P, Hariss D, et al. Nitric oxide as a double-edged sword in pulmonary viral infections: Mechanistic insights and potential therapeutic implications. Gene. 2024;899:148148.

53. Xue Q, Yan Y, Zhang R, Xiong H. Regulation of iNOS on Immune Cells and Its Role in Diseases. Int J Mol Sci. 2018;19(12):3805.

54. Omar M, Abdelal HO. Nitric oxide in parasitic infections: a friend or foe? J Parasit Dis. 2022;46(4):1147–1163.

55. Mishra BB, Lovewell RR, Olive AJ, et al. Nitric oxide prevents a pathogen-permissive granulocytic inflammation during tuberculosis. Nat Microbiol. 2017;2:17072.

56. Dal Secco D, Paron JA, de Oliveira SH, Ferreira SH, Silva JS, Cunha Fde Q. Neutrophil migration in inflammation: nitric oxide inhibits rolling, adhesion and induces apoptosis. Nitric Oxide. 2003;9(3):153–164.

57. Yau B, Mitchell AJ, Too LK, Ball HJ, Hunt NH. Interferon-gamma-Induced Nitric Oxide Synthase-2 Contributes to Blood/Brain Barrier Dysfunction and Acute Mortality in Experimental *Streptococcus pneumoniae* Meningitis. J Interferon Cytokine Res. 2016;36(2):86–99.

58. Perrone LA, Belser JA, Wadford DA, Katz JM, Tumpey TM. Inducible nitric oxide contributes to viral pathogenesis following highly pathogenic influenza virus infection in mice. J Infect Dis. 2013;207(10):1576–1584.

59. Fernandes KS, Neto EH, Brito MM, Silva JS, Cunha FQ, Barja-Fidalgo C. Detrimental role of endogenous nitric oxide in host defence against *Sporothrix schenckii*. Immunology. 2008;123(4):469–479.

60. Burke TP, Engstrom P, Chavez RA, Fonbuena JA, Vance RE, Welch MD. Inflammasome-mediated antagonism of type I interferon enhances *Rickettsia* pathogenesis. Nat Microbiol. 2020;5(5):688–696.

61. Salim T, Sershen CL, May EE. Investigating the Role of TNF-alpha and IFN-gamma Activation on the Dynamics of iNOS Gene Expression in LPS Stimulated Macrophages. PLoS One. 2016;11(6):e0153289.

62. Hsu YH, Chen HI. Pulmonary pathology in patients associated with scrub typhus. Pathology. 2008;40(3):268–271.

63. Kitano T, Yamada H, Kida M, Okada Y, Saika S, Yoshida M. Impaired Healing of a Cutaneous Wound in an Inducible Nitric Oxide Synthase-Knockout Mouse. Dermatol Res Pract. 2017;2017:2184040.

64. Yamasaki K, Edington HD, McClosky C, et al. Reversal of impaired wound repair in iNOS-deficient mice by topical adenoviral-mediated iNOS gene transfer. J Clin Invest. 1998;101(5):967–971.

65. Ishida H, Ray R, Amnuaysirikul J, Ishida K, Ray P. Nitric oxide synthase gene transfer overcomes the inhibition of wound healing by sulfur mustard in a human keratinocyte in vitro model. ISRN Toxicol. 2012;2012:190429.

66. Ahmed R, Augustine R, Chaudhry M, et al. Nitric oxide-releasing biomaterials for promoting wound healing in impaired diabetic wounds: State of the art and recent trends. Biomed Pharmacother. 2022;149:112707.

67. Chung MH, Lee JS, Baek JH, Kim M, Kang JS. Persistence of *Orientia tsutsugamushi* in humans. J Korean Med Sci. 2012;27(3):231–235.

68. Smadel JE, Ley HL, Jr., Diercks RH, Cameron JA. Persistence of *Rickettsia tsutsugamushi* in tissues of patients recovered from scrub typhus. Am J Hyg. 1952;56(3):294–302.

69. Liang Y, Wang H, Gonzales C, et al. CCR7/dendritic cell axis mediates early bacterial dissemination in *Orientia tsutsugamushi*-infected mice. Front Immunol. 2022;13:1061031.

70. Salje J. *Orientia tsutsugamushi*: A neglected but fascinating obligate intracellular bacterial pathogen. PLoS Pathog. 2017;13(12):e1006657.

71. Xu G, Mendell NL, Liang Y, et al. CD8+ T cells provide immune protection against murine disseminated endotheliotropic *Orientia tsutsugamushi* infection. PLoS Negl Trop Dis. 2017;11(7):e0005763.

72. Hauptmann M, Kolbaum J, Lilla S, et al. Protective and Pathogenic Roles of CD8+ T Lymphocytes in Murine *Orientia tsutsugamushi* Infection. PLoS Negl Trop Dis. 2016;10(9):e0004991.

73. Wei XQ, Charles IG, Smith A, et al. Altered immune responses in mice lacking inducible nitric oxide synthase. Nature. 1995;375(6530):408–411.

74. McInnes IB, Leung B, Wei XQ, Gemmell CC, Liew FY. Septic arthritis following *Staphylococcus aureus* infection in mice lacking inducible nitric oxide synthase. J Immunol. 1998;160(1):308–315.

75. MacLean A, Wei XQ, Huang FP, Al-Alem UA, Chan WL, Liew FY. Mice lacking inducible nitric-oxide synthase are more susceptible to herpes simplex virus infection despite enhanced Th1 cell responses. J Gen Virol. 1998;79 (Pt 4):825–830.

76. Soong L, Mendell NL, Olano JP, et al. An Intradermal Inoculation Mouse Model for Immunological Investigations of Acute Scrub Typhus and Persistent Infection. PLoS Negl Trop Dis. 2016;10(8):e0004884.

77. Sunyakumthorn P, Paris DH, Chan TC, et al. An intradermal inoculation model of scrub typhus in Swiss CD-1 mice demonstrates more rapid dissemination of virulent strains of *Orientia tsutsugamushi*. PLoS One. 2013;8(1):e54570.

78. Roshick C, Wood H, Caldwell HD, McClarty G. Comparison of gamma interferon-mediated antichlamydial defense mechanisms in human and mouse cells. Infect Immun. 2006;74(1):225–238.

79. Tenopoulou M, Doulias PT. Endothelial nitric oxide synthase-derived nitric oxide in the regulation of metabolism. F1000Res. 2020;9:F1000 Faculty Rev-1190.

80. Forstermann U, Munzel T. Endothelial nitric oxide synthase in vascular disease: from marvel to menace. Circulation. 2006;113(13):1708–1714.

